# Host-environment interplay shapes fungal diversity in mosquitoes

**DOI:** 10.1101/2020.12.01.407494

**Authors:** Patil Tawidian, Kerri L. Coon, Ari Jumpponen, Lee W. Cohnstaedt, Kristin Michel

## Abstract

Mosquito larvae encounter diverse assemblages of bacteria (i.e. ‘microbiota’) and fungi (i.e. ‘mycobiota’) in the aquatic environments they develop in. However, while a number of studies have addressed the diversity and function of microbiota in mosquito life history, relatively little is known about mosquito-mycobiota interactions outside of several key fungal entomopathogens. In this study, we used high-throughput sequencing of ITS2 gene amplicons to provide the first simultaneous characterization of the mycobiota in field-collected *Aedes albopictus* larvae and their associated aquatic environments. Our results reveal unprecedented variation in mycobiota among adjacent but discrete larval breeding habitats. Our results also reveal distinct mycobiota assembly in the mosquito gut versus other tissues, with gut-associated fungal communities being most similar to those present in the environment where larvae feed. Altogether, our results identify the environment as the dominant factor shaping mosquito mycobiota with no evidence of environmental filtering of the gut mycobiota. These results also identify mosquito feeding behavior and fungal mode of nutrition as potential drivers of tissue-specific mycobiota assembly after environmental acquisition.

**IMPORTANCE:** The Asian tiger mosquito, *Aedes albopictus*, is the dominant mosquito species in the USA and an important vector of arboviruses of major public health concern. One aspect of mosquito control to curb mosquito-borne diseases has been the use of biological control agents such as fungal entomopathogens. Recent studies also demonstrate the impact of mosquito-associated microbial communities on various mosquito traits, including vector competence. However, while much research attention has been dedicated to understanding the diversity and function of mosquito-associated bacterial communities, relatively little is known about mosquito-associated fungal communities. A better understanding of the factors that drive mycobiota diversity and assembly in mosquitoes will be essential for future efforts to target mosquito micro- and mycobiomes for mosquito and mosquito-borne disease control.

## INTRODUCTION

Mosquito larvae and adults continuously encounter diverse microorganisms in their aquatic and terrestrial environments (1-3). These microorganisms include bacteria and fungi, which assemble into bacterial and fungal communities, hereafter referred to as micro- and mycobiota, respectively. The mosquito gut microbiota plays a profound role in the growth and development of larvae, as well as adult survival, fecundity, and mosquito-borne transmission of disease-causing pathogens (4-11). In contrast, studies of fungus-mosquito interactions have largely focused on the identification of fungal entomopathogens and their use as bioinsecticides to control mosquito larvae and adults (12-16). Of the 158 fungal species observed in or isolated across mosquito species, nearly two-thirds are entomopathogens (3). These entomopathogenic fungi infect mosquitoes mostly through the cuticle, and rarely through ingestion (3). However, not all fungus-mosquito interactions have a negative outcome on mosquitoes. Fungi such as yeasts can be sufficient for mosquito development as a nutritional source and because they induce gut hypoxia, which serves as a cue for larval development (10, 17, 18). Taken together, these studies strongly suggest that fungi have the potential to profoundly impact mosquito biology; yet, only few have examined the factors shaping mycobiota in larvae and their larval breeding environments where they naturally develop.

Studies that have focused on the characterization of microbial diversity in mosquitoes collectively indicate that bacterial communities can vary substantially across different larval environments and between individuals that co-occur in the same environment, even at small local scales. Studies also indicate that the majority of bacteria present in mosquitoes are restricted to the gut (19–21). The few available published mycobiota studies suggest that environment may also be a dominant factor shaping fungal diversity in mosquitoes (22). However, to date no study has simultaneously characterized mycobiota in mosquito larvae and the aquatic environment they inhabit. In addition, studies thus far have largely focused on the characterization of mycobiota in only either whole mosquitoes or dissected guts. Fungi also form associations with their mosquito hosts as a function of their mode of nutrition: some taxa enter the mosquito through the body surface (cuticle), whereas others enter via ingestion through the gut. However, no study to date has examined whether fungal communities differ between mosquito host tissues.

The overall goal of this study was to determine the factors that shape fungus-mosquito interactions in larvae of the Asian tiger mosquito (*Aedes albopictus*), an abundant mosquito species of public health concern because adult females transmit the causative agents of Dengue fever, Chikungunya, and Zika (23–26). *Ae. albopictus* is ubiquitous in urban and peri-urban areas throughout most of the world, where larvae inhabit diverse natural and artificial containers (27–29), and feed on living and decaying organic matter using diverse filtering, grazing, and shredding behaviors (30, 31). Here, we sampled water and *Ae. albopictus* late-stage (L4) larvae from several types of man-made container breeding sites on a fine geographic scale. We then used ITS2 metabarcoding to determine the mycobiota composition and diversity of mosquito larvae and their larval breeding water. Using these data, we determined that the aquatic environment is a major driver of mosquito mycobiota composition. In addition, we determined additional drivers such as mosquito feeding behavior and fungal mode of nutrition contribute to the mycobiota and its diversity in different mosquito tissues.

## RESULTS

### Fungal communities based on ITS2 metabarcoding

We analyzed the mycobiota in *Ae. albopictus* larvae and water sampled from ten aquatic breeding sites located within a 11.6-km^2^ area in Manhattan, KS (**Fig. 1**). From each site, we sampled ~50-ml of water and ten individual larvae. The larvae were aseptically dissected to produce paired gut and carcass samples prior to sequencing fungal ITS2 metabarcode amplicons on an Illumina MiSeq. The final dataset consisted of a total of 4,259,124 quality-filtered sequences, assigned to a total of 3,415 operational taxonomic units (OTUs) at a 97% sequence similarity. Rarefaction curves saturated or nearly saturated at 5,000 sequences for most samples, indicating that the fungal diversity was captured in our sampling (**Fig. S1**). Six fungal phyla were identified across all samples and accounted for 91.1% of the total quality-filtered reads: Ascomycota (59.5%), Basidiomycota (30.8%), Chytridiomycota (0.316%), Glomeromycota (0.057%), Mucoromycota (0.465%), and Rozellomycota (0.019%) (**Fig. 2**). The remaining 8.86% of reads could not be classified past the Kingdom Fungi but were included in all downstream analyses.

**Figure 1.**
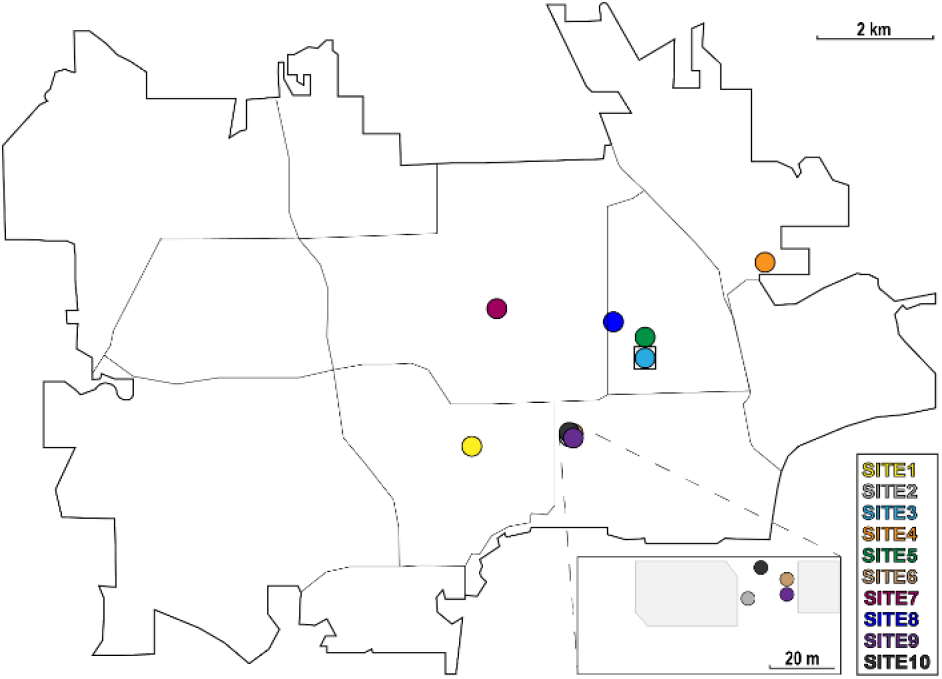
Collection sites for the ITS2 libraries prepared from water and *Aedes albopictus* mosquito larvae. Figure depicts the location of each collection site in Manhattan, KS. Sites 1 and 2 are natural *Ae. albopictus* breeding sites, while the remaining sites were artificially constructed using plastic mosquito oviposition cups lined with germination paper. The square surrounding site 3 indicates that this site was eliminated from downstream community analyses due to low sequencing reads.

**Figure 2.**
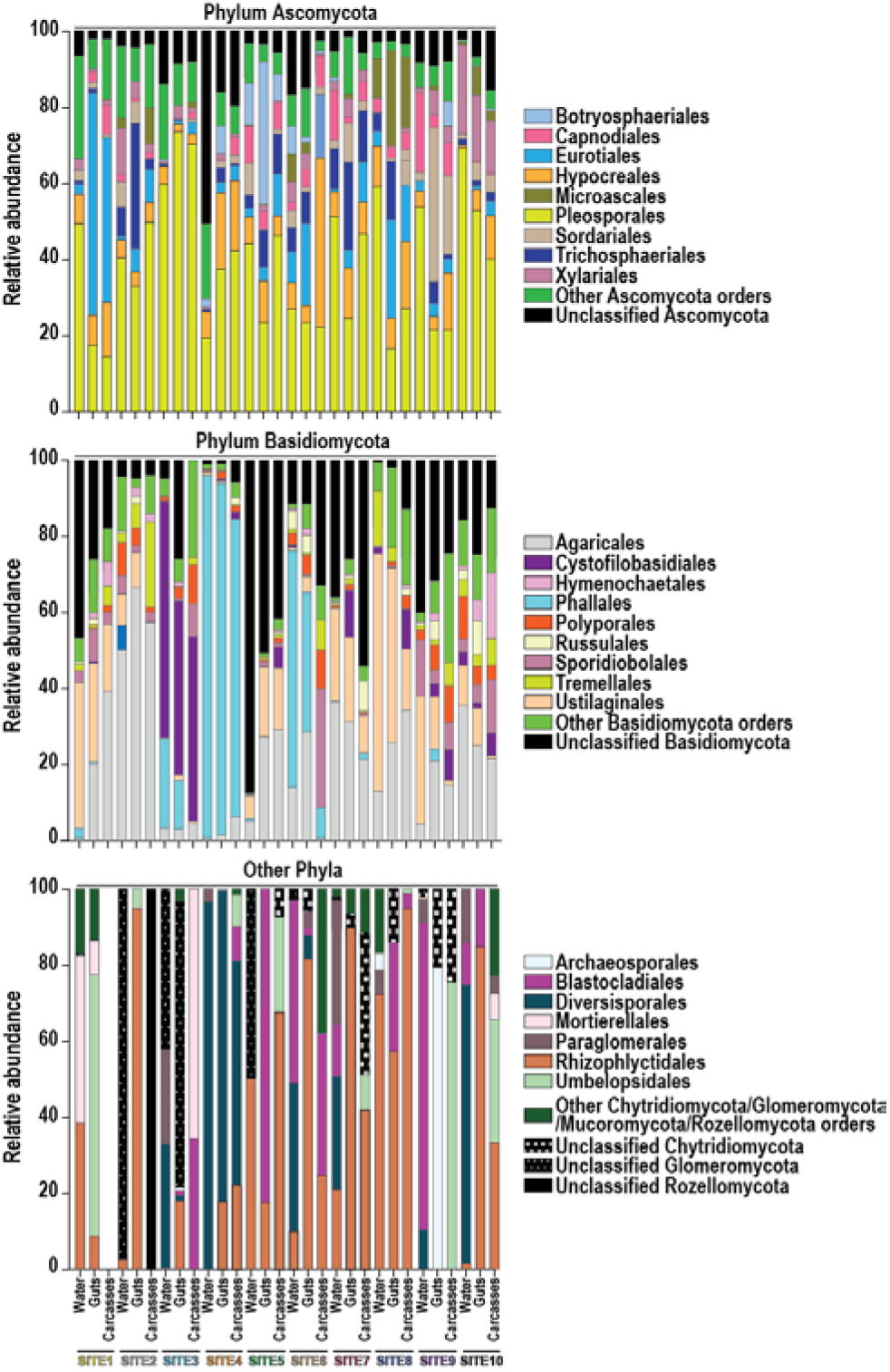
Fungi at the levels of phylum and order in water and mosquito larvae from each collection site. Gut and carcass samples from individual larvae collected from a given site were pooled for the bar graphs presented. For a given phylum, bars present the proportion of sequencing reads assigned to that phylum that were also classified to a specific fungal order. All orders that represented ≥ 10% of the reads from a given sample are listed in the legend; less abundant orders are grouped together under the ‘Other’ categories.

Within the phylum Ascomycota, OTUs in the fungal orders Pleosporales, Hypocreales, and Eurotiales were shared across all water and mosquito samples and represented ~38.1%, 10.2%, and 9.5% of the total reads assigned to this phylum, respectively (**Fig. 2**). Within the phylum Basidiomycota, the majority of reads (~21.8%) were associated with the order Agaricales and reads assigned to this order were detected in all of the water and mosquito samples (**Fig. 2**). In contrast, fungi in the orders Cystofilobasidiales and Phallales were abundant in the water, mosquito gut and carcass samples from only sites 3 and 4 (20.9% and 31.5%, respectively) but rare (on average 1.87%) in samples from other sites (**Fig. 2**). The majority of reads (on average 45.0%) within the remaining four phyla belonged to the orders Rhizophlyctidales (phylum Chytridiomycota) and Mortierellales (phylum Mucoromycota) (**Fig. 2**).

### Local environment is the dominant factor that shapes the mosquito mycobiota

To identify whether environment is a major driver of mycobiota diversity in mosquito larvae, we visualized Bray-Curtis dissimilarities among all our water, gut, and carcass samples using a principal coordinate analysis (PCoA, **Fig. 3**). To determine whether samples were distinct between breeding sites, we analyzed the distance matrices using permutational multivariate analysis of variance (PERMANOVA). The results revealed significant differences in the mycobiota between samples from different sites (**Fig. 3**). We then ran a multivariate analysis of variance (MANOVA) on the distance matrices of the first three PCoA vectors to determine whether the mycobiota between breeding sites and/or mosquito tissue type (gut vs. carcass) were significantly different. Univariate analyses of variance (ANOVAs) were performed on significant MANOVA factors to examine which axis or axes drove any patterns observed in the PCoA plots. The MANOVA and univariate ANOVAs further indicated that differences along all three PCoA axes were significant for site and the interaction of site and mosquito tissue type, but samples only separated by tissue along the third axis (**Table 1**).

**Figure 3.**
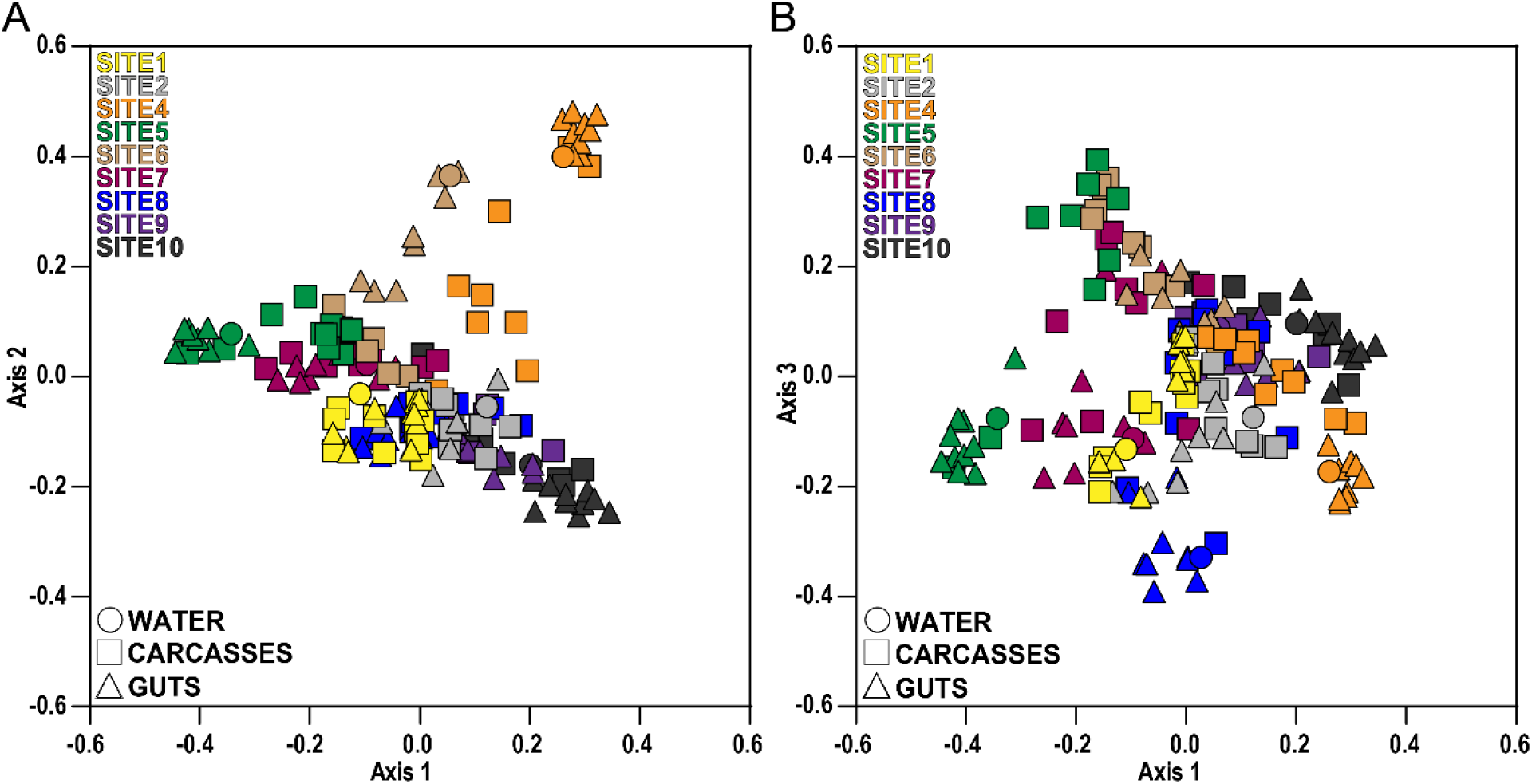
Principal coordinates analysis based on pairwise Bray-Curtis distances. The legends in the upper left of each plot designate collection site by color, while legends in the lower left of each plot designate sample type (water vs. mosquito gut or carcass) by symbol shape. Samples cluster significantly by collection site (PERMANOVA, F = 7.167, *P* < 0.001).

**Table 1.** The contribution of collection sites and mosquito tissues on the mosquito larval mycobiota.

| ***A - Multivariate analysis of mosquito guts and carcasses (mosquito tissues) across all collection sites*** |  |  |  |  |  |
| --- | --- | --- | --- | --- | --- |
| **Source of variation** | **Pillai's trace** | **Hypothesis df** | **Error df** | **F** | **P** |
| Collection sites | 2.26 | 24.0 | 456 | 57.8 | < 0.0001 |
| Mosquito tissues | 0.348 | 3.00 | 150 | 26.7 | < 0.0001 |
| Collection sites:<br>Mosquito tissues | 1.04 | 24.0 | 456 | 10.0 | < 0.0001 |
| ***B - Univariate analysis of the PCoA axes driving the variation between mosquito tissues*** |  |  |  |  |  |
| **Source of variation** | **Sum of squares** | **Residual sum of squares** | **df** | **F** | **P** |
| Axis 1 | 0.001 | 5.55 | 1.00 | 0.039 | 0.844 |
| Axis 2 | 0.018 | 4.52 | 1.00 | 0.686 | 0.409 |
| Axis 3 | 0.616 | 4.34 | 1.00 | 27.8 | < 0.0001 |
| ***C - Univariate analysis of the PCoA axes driving the variation between collection sites*** |  |  |  |  |  |
| **Source of variation** | **Sum of squares** | **Residual sum of squares** | **df** | **F** | **P** |
| Axis 1 | 4.38 | 5.55 | 8.00 | 75.4 | < 0.0001 |
| Axis 2 | 3.38 | 4.52 | 8.00 | 59.8 | < 0.0001 |
| Axis 3 | 1.83 | 4.34 | 8.00 | 14.6 | < 0.0001 |
| ***D - Univariate analysis of the PCoA axes driving an interaction between collection sites and tissues*** |  |  |  |  |  |
| **Source of variation** | **Sum of squares** | **Residual sum of squares** | **df** | **F** | **P** |
| Axis 1 | 0.390 | 1.17 | 8.00 | 9.52 | < 0.0001 |
| Axis 2 | 0.453 | 1.12 | 8.00 | 12.9 | < 0.0001 |
| Axis 3 | 0.591 | 1.89 | 8.00 | 8.60 | < 0.0001 |

### Tissue-specific patterns in mycobiota assembly

To test whether patterns of mycobiota assembly differ between the mosquito gut and other tissues, we next compared calculated indices for alpha diversity and community composition within all water and individual mosquito gut and carcass samples we collected. Alpha diversity differed between mosquitoes tissues (gut vs. carcass) as measured by observed (W = −3,165, P < 0.0001) and extrapolative Chao1 (W = −3,153, P < 0.0001) OTU richness, as well as Shannon’s diversity (W = −1,722, P < 0.0001). Fungal species richness and diversity were consistently higher in mosquito gut samples than in the corresponding carcass samples (**Table S1**). Alpha diversity was also generally higher in the water than in the mosquito samples (**Table S1**). Interestingly, the pairwise Bray-Curtis dissimilarities were higher between water and carcass samples than between water and gut samples (**Fig. 4; Fig. S2**), indicating that the water and mosquito gut mycobiota were on average more similar than those of the water and mosquito carcasses. We also analyzed the distance matrices between individual gut and carcass samples by site using PERMANOVA. The two mosquito tissues differed in a majority of sites suggesting distinct fungal community assembly in mosquito guts and carcasses (**Fig. S3**).

**Figure 4.**
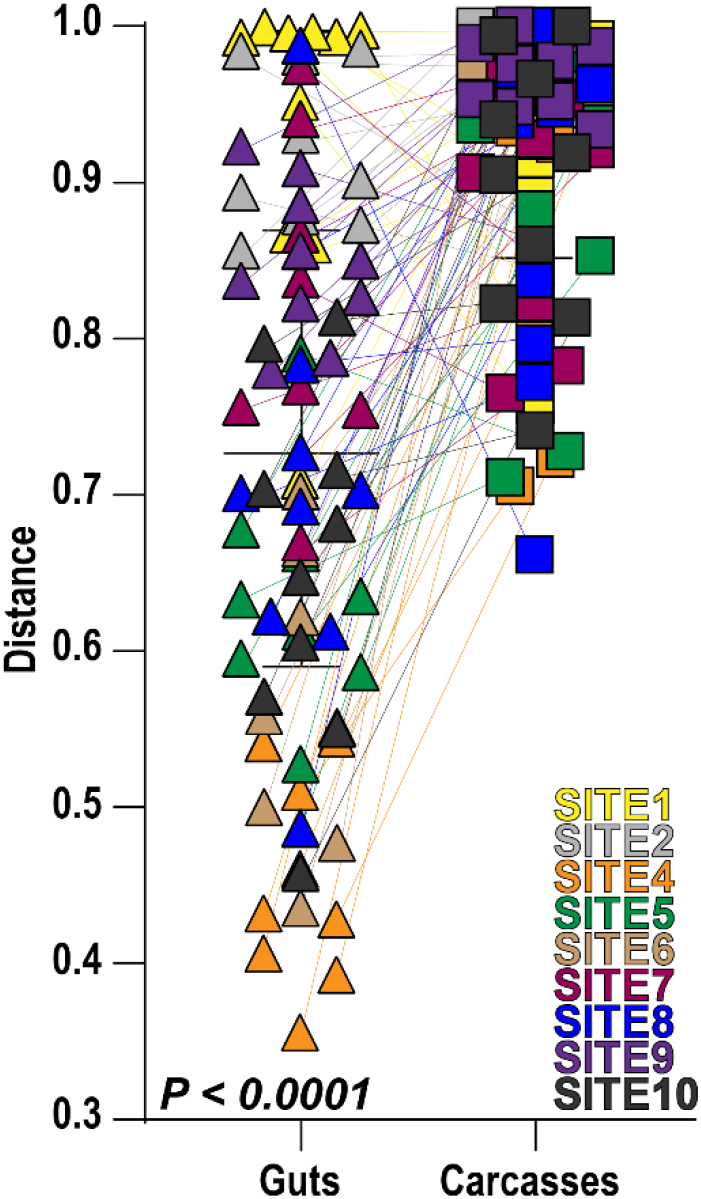
Bray-Curtis distances between fungal communities detected in water and either the guts or carcasses of mosquito larvae collected from the same site. Connected lines depict matching mosquito gut and carcass samples for a given collection site. Statistical significance was determined using a Wilcoxon signed-rank test.

### Mosquito feeding behavior and fungal mode of nutrition drive tissue-specific patterns in mycobiota assembly

To identify specific fungal OTUs that are differentially associated with the mosquito gut and carcass, we performed indicator species analyses using two different indices of the 149 OTUs that made up 80% of total reads in the individual mosquito gut and carcass samples across all sites. Using the indicator value Index (IndVal), we identified 41 indicator OTUs for mosquito guts across all breeding sites (**Fig. 5A**) (**Table S2**). In contrast to the mosquito guts, the IndVal analysis identified no indicator OTUs of mosquito carcasses. The 41 mosquito gut indicator OTUs were assigned to six ecological guilds: saprophyte (34%), plant pathogen (25%), endophyte (8.9%), animal pathogen (5.3%), epiphyte (1.7%), and ectomycorrhizal (1.7%). The remaining 23% were not assigned ecological guilds.

**Figure 5.**
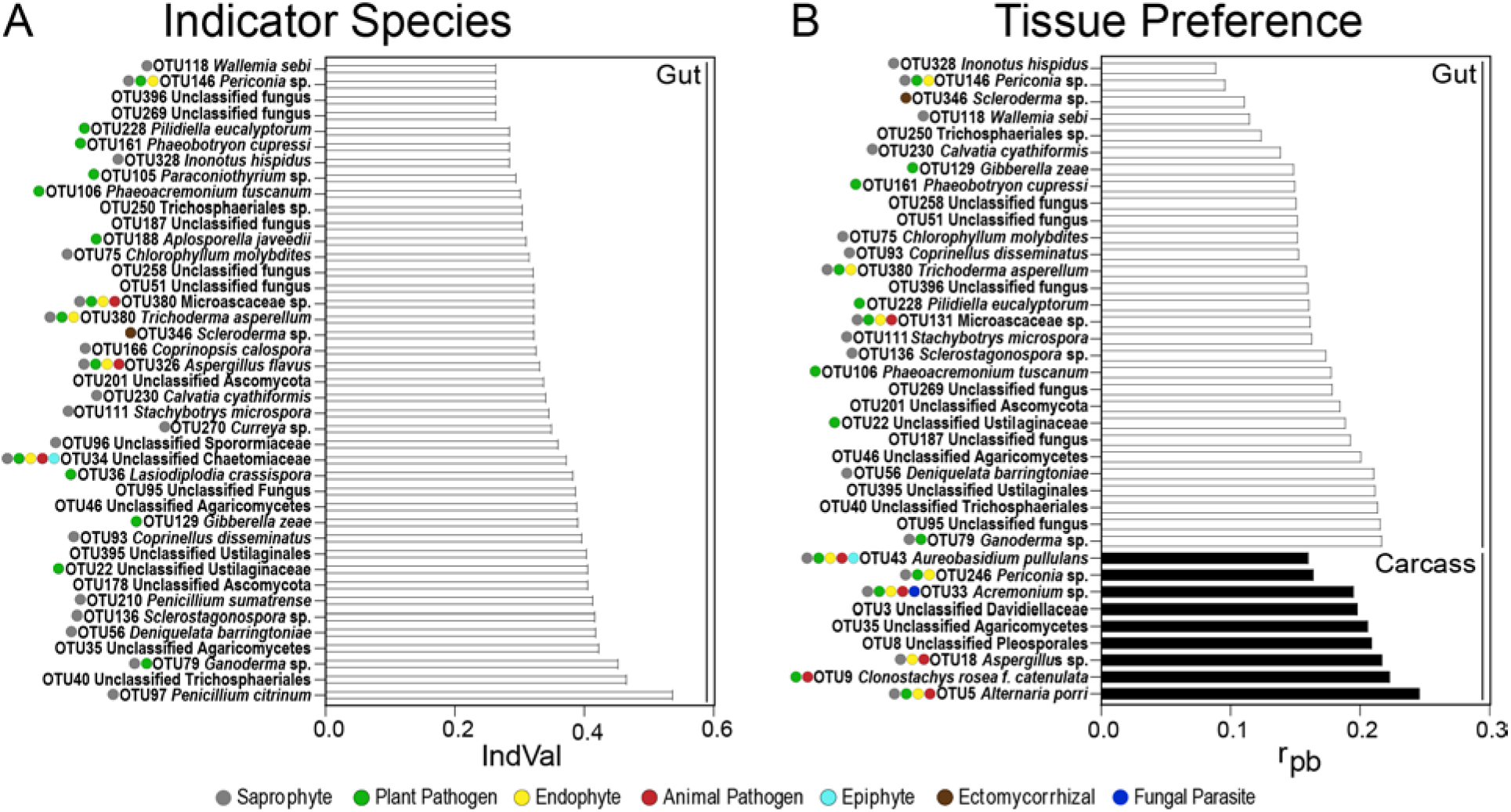
Indicator species and tissue preference of fungal OTUs between mosquito guts and carcasses across all larval breeding sites. (A) The characteristic fungal OTUs of mosquito guts across all breeding sites were identified using the indicator value index (IndVal) at p-value of 0.05 and 9,999 permutations. (B) OTU tissue preference between mosquito guts and carcasses across breeding sites were identified using the point-biserial correlation index (rpb) at p-value of 0.05 and 9,999 permutations. Ecological guilds inferred by FUNGuild are organized from the most abundant to the least abundant guild.

We then used the point-biserial correlation index (rpb) to determine the impact of fungal niche preference on differential mycobiota assembly between mosquito guts and carcasses (**Fig. 5B**) (**Table S3**). We found that 29 OTUs, all of which were also gut indicator OTUs, significantly associated with mosquito guts (**Table S3**). The OTUs with mosquito gut preference were primarily assigned the ecological guilds of saprophyte (32.4%), plant pathogen (21.6%), and endophyte (8.1%), followed by animal pathogen (2.7%) and ectomycorrhiza (2.7%). The remaining 32.4% of OTUs could not be assigned to any ecological guild. The rpb analysis identified nine OTUs that significantly associated with mosquito carcasses (**Table S3**). The majority of these OTUs belonged to four ecological guilds: saprophyte (21.7%), endophyte (21.7%), animal pathogen (17.4%), and plant pathogen (17.4%), followed by fungal parasite (4.3%) and epiphyte (4.3%). The remaining 13% of OTUs were not assigned to any ecological guild.

In addition to these indicator analyses, we also mined the dataset for taxa with known entomopathogenic potential. We identified one OTU (OTU68) assigned to the genus *Beauveria*, which contains several known entomopathogens, and another (OTU9) assigned to *Clonostachys rosea f. catenulata* which significantly associated with mosquito carcasses in the rpb analysis. Sequences assigned to a member of the genus *Beauveria* were present in mosquito samples from 50% of sampled breeding sites. Within these sites, *Beauveria sp*. was found only sporadically (in 15 guts and seven carcasses from 48 mosquito larvae total), and only five larvae had OTU reads assigned to *Beauveria* in both guts and carcasses. The distribution of reads across these two tissue types in those five larvae did not show a clear trend, ranging from 10-fold higher reads in guts as compared to carcasses and vice versa. Sequences assigned to *Clonostachys rosea f. catenulata* were present in only three of the nine sampled breeding sites. Within these sites, *C. rosea f. catenulata* was found in 24 gut and 23 carcass samples. In the 21 larvae where *C. rosea f. catenulata* were found in both guts and carcasses, the number of reads was on average 19-fold higher in the carcass as compared to the gut samples.

## DISCUSSION

This study reports the mycobiota composition of *Ae. albopictus* larvae and their natural breeding habitats on a small geographic scale. The breeding water and mosquito guts and carcasses sampled in our study harbored a diverse, uneven, and rich fungal community. Similar patterns of fungal diversity were reported in natural and artificial mosquito breeding sites sampled from different regions in Taiwan and in mosquito adults collected intercontinentally (22, 32). In our study, the fungal communities identified in the breeding water and mosquito guts and carcasses were dominated by fungi in the phyla Ascomycota and Basidiomycota. This parallels previously reported fungal communities in water and organic substrates collected from tree holes and man-made containers (22, 33). Our results are not surprising and can be explained by the ubiquity of Ascomycota and Basidiomycota in freshwater ecosystems as compared to the remaining fungal phyla (34-36). The ITS2 (fITS7 and ITS4) primers we used for amplification are commonly used in mycobiota surveys (37-41) and readily identify members of the fungal phyla Ascomycota, Basidiomycota, Chytridiomycota, and Mucoromycota, as demonstrated by our mock fungal community. However, ITS2 primers show low species identification and discrimination in early diverging fungal lineages (42, 43). Additionally, a considerable percentage (9%) of the OTUs assigned to unclassified fungi had to be reclassified upon closer inspection, most commonly to algae, which is consistent with previous studies reporting the amplification of algae by the fITS7 and ITS4 primer pair (44). Future studies using these primers should therefore also carefully inspect OTUs assigned to non-classified fungi.

The overall goal of this study was to determine the drivers of mycobiota assembly of field-collected *Ae. albopictus* larvae. Our results show that the fungal communities in the larvae of this mosquito species largely reflect those of their breeding environments. To our knowledge, this is the first mycobiota data set with sampling at a local scale and sequences obtained from individual larvae from which the guts were dissected. Our results corroborate data of factors shaping the bacterial diversity in larvae of various mosquito species. Gimonneau et al (45) showed that the bacterial community in field-collected *Anopheles gambiae* and *Anopheles coluzzii* mosquito larvae largely overlapped with that of the aquatic habitat, suggesting that the larval breeding water is the major source of the microbiota assembly in mosquito larvae. Similar findings were also reported for the larval microbiota of several *Aedes, Anopheles*, and *Culex* species, including *Aedes japonicus, Aedes triseriatus, Aedes aegypti, Ae. albopictus, Anopheles darlingi, Anopheles nuneztovari, Culex quinquefasciatus*, and *Culex restuans*, collected from several man-made container breeding sites (6, 46, 47). We recognize that our study only assessed the mycobiota composition in a single mosquito species. Hence, we cannot exclude the host as a significant factor shaping the mycobiota assembly in *Ae. albopictus* mosquito larvae. However, our results provide little evidence for filtering or enrichment of specific taxa which are shared across breeding sites, further supporting the dominant role of the larval aquatic habitat in mycobiota assembly.

Our results further show that the mycobiota of the mosquito larvae gut differs from that of the carcass. Although not identical with the larval breeding water, the gut mycobiota was more similar to the breeding environment than it was to the mycobiota of the carcass. The similarity of the gut mycobiota to the environment can be explained by the filter-feeding habit of mosquito larvae in general, which presumably captures a large portion of fungal taxa from the breeding water (31). *Ae. albopictus* larvae also display other feeding habits, including grazing and shredding on decaying leaf matter (48–50), which further contributes to the gut fungal diversity. This is supported by the results of the indicator species analysis where the gut indicator OTUs predominantly are saprophytes, endophytes, and pathogens that typically occur in plant tissues. To exemplify, one of the indicator OTUs belongs to the family Ustilaginaceae, which is composed of more than 1,200 obligately biotrophic fungal species that can infect more than 4,000 plant species (51). Members of this family can infect forage grasses and crops such as corn, barley, and wheat (51, 52). Several other plant associated fungi that were abundant and frequent in the gut samples, include *Gibberella zeae, Phaeoacremonium tuscanum*, and *Phaeobotryon cupressi* taxa that likely represent putative pathogens of diverse plant crops, including wheat and grapevine, and trees (53-57). This further supports that the fungal diversity observed in the larval gut of mosquitoes is a consequence of feeding on plant material.

In contrast, the fungal diversity of mosquito carcasses, from which we had removed the heads to exclude any fungi captured on the mouth brushes during feeding, was lower and contained communities distinct from those in the larval guts and breeding water. Approximately half of the OTUs with preference to carcass compared to gut were putative animal pathogens, suggesting that these OTUs either interact with the carcass by attaching to the cuticle or infect the carcass through the gut after ingestion (58-64). These included OTUs assigned to the species *C. rosea*, a fungal entomopathogen of several insect hosts, including leafhoppers, whiteflies, and alfalfa weevils (65-67), and *Alternaria porri*, which causes mortality in green apple aphids and delays hatch rates in egg masses of the European corn borer (68, 69). These two OTUs were also detected in gut samples, suggesting that ingestion might have led to systemic infection by these fungi. Overall, our data strongly suggest that in addition to mosquito feeding behavior, fungal ecology and niche preference further separates fungal communities in *Ae. albopictus* larval tissues.

Overall, our analyses found little additional evidence of *Ae. albopictus* infections with known fungal entomopathogens of mosquitoes. Of the 67 fungal species considered entomopathogenic and/or entomotoxigenic to mosquitoes (3), *Beauveria sp*. was the only previously described mosquito entomopathogen that we detected. This observation is consistent with a previous study describing the mycobiota of field-collected *Ae. albopictus* adult females (32). However, the distribution of sequences in our dataset provides little support for active mosquito infection by *Beauveria* in our collection sites. The sequences assigned to *Beauveria* sp. were rare and sporadic across the mosquito guts and even less commonly found in the carcass samples. Nevertheless, we were able to isolate a local strain of *Beauveria bassiana* from a single *Ae. albopictus* larva collected at site 5. Future studies with this isolate will test the efficacy of this strain to infect *Ae. albopictus* larvae.

To our surprise, we did not detect OTUs assigned to the genus *Smittium*, an early diverging fungal lineage in the subphylum Kickxellomycotina, which contains several species of gut symbionts and pathogens of mosquito larvae (70–74). Despite the large host range of the genus *Smittium*, to our knowledge, it has not been reported in *Ae. albopictus* larvae. It is thus possible that members of this genus are not able to colonize the gut of *Ae. albopictus* larvae. It is also possible that *Smittium sp*. do not persist in mosquito breeding grounds in Kansas and therefore were not detected in our dataset. However, an equally parsimonious explanation is a lack of detection due to the choice of primers that poorly capture early diverging and basal fungal phyla (75). In future studies, we will therefore employ taxon-specific primers and microscopy techniques to assess the presence of *Smittium* sp. in *Ae. albopictus* larvae as described previously (76–78).

In conclusion, this study provides fundamental insights into the broad range of encounters between mosquito larvae and fungi in the larval breeding water. Our results show that mosquito breeding water harbors a highly rich and diverse fungal community on a fine geographic scale, which drives mosquito mycobiota assembly. We further show the contribution of mosquito feeding behavior and fungal ecology to tissue-specific patterns of mycobiota assembly. Future studies will have to assess whether these observed patterns can be generalized across different mosquito species, and whether ontogeny further contributes to mosquito mycobiota assembly.

## MATERIALS AND METHODS

### Sample collections

*Ae. albopictus* L4 larvae and corresponding water were sampled from a total of ten breeding sites in Manhattan, KS during 2017 and 2018—two naturally occurring mosquito breeding sites and eight artificial (man-made) breeding sites consisting of plastic mosquito oviposition cups lined with heavy weight seed germination paper (Anchor paper co, MN, USA) (**Fig. 1; Table S1**). For the 2017 larval collections, mosquito larvae were kept in their respective environments, transported to the laboratory in labeled plastic containers (Bare Eco-Forward Rpet Deli Container), and incubated at 27°C with 75% RH for 24 hours before larval gut dissections. For the 2018 larval collections, mosquito larvae were transported to the laboratory as in 2017 but dissected immediately upon arrival. Water samples were immediately stored at −80°C until nucleic acid extraction and sequencing.

### Mosquito dissections

Mosquito larvae were surface washed six times with sterile Milli-Q water before decapitation (to exclude any transiently attached fungi from the environment on the mouth brushes) and dissection. Dissection of the larval gut from the body, hereafter referred to as mosquito carcass, was accomplished using flame-sterilized forceps and dissecting pins. Dissected gut and carcass samples were immediately frozen with liquid nitrogen and stored at −80°C until further processing. Finely cut, bleached nets were used as negative controls to screen for contamination during the dissection process. These negative controls were processed simultaneously with mosquito larva dissections.

### Sample preparation and Illumina MiSeq

Total DNA was extracted using the DNeasy® PowerSoil® Kit (MoBio Laboratory, Carlsbad, CA, USA) following the manufacturer’s instructions with minor modifications from a total of 100 mosquito gut and carcass samples, ten water samples filtered through 1-micron nuclepore membranes (Whatman®), and seven dissection control samples. DNA samples were stored at −20°C until PCR amplification. Extracted DNA was quantitated using NanoDropTM 2000/2000c spectrophotometers (Thermo Scientific, Waltham, MA, USA) and standardized to 2 ng/μL concentration. The fungal amplicon library was generated by triplicate PCR amplification using barcoded forward primer fITS7 (5’-GTGARTCATCGAATCTTTG-3’) and barcoded reverse primer ITS4 (5’-TCCTCCGCTTATTGATATGC-3’) following a protocol described by (79) with minor modifications. PCR with 20 ng template DNA included an initial denaturation of 30 s at 98 °C, 35 cycles of denaturing, annealing, and extension at 98 °C for 10 s, 54 °C for 30 s, 72 °C for 1 min, followed by a final 72 °C extension step for 9 min. The PCR negative control was certified nuclease-free sterile water. A mock fungal community was created with DNA from ten fungal species belonging to different phyla to determine sequencing quality and range of fungal taxa identification (**Table S4**).

Successful PCR amplification was determined by visualizing 5 μL of the PCR products on 1% agarose gels. The remaining 45 μL from each triplicate PCRs were pooled and purified using Mag-Bind® RXNPure plus (Omega Bio-tek; Norcross, GA, USA). The clean amplicons from each sample were pooled at equal concentrations. Illumina MiSeq adaptors were ligated onto the amplicon library using a NEBNext® DNA MasterMix for Illumina Kit (New England BioLabs, Ipswitch, MA, USA) and sequences were generated using a MiSeq instrument (2 × 250 cycles; Illumina, San Diego, CA, USA) at the Kansas State University Integrated Genomics Facility (Manhattan, KS, USA).

### Sequence processing

Paired-end sequences were processed using mothur (v. 1.38.1) (80). Sequences with ambiguous bases, mismatches to primers, and homopolymers longer than 10 bp were removed. A total of 229,195 chimeric sequences were identified using the VSEARCH algorithm (81) and removed from the dataset. Fungal sequences were assigned to taxa using the naïve bayesian classifier against the UNITE-curated International Nucleotide Sequence Database reference database (82, 83). During data processing through mothur, 191,081 sequences were either unassigned or assigned to unclassified plantae and the protozoan phyla cercozoa and ciliophora, respectively, and removed from the dataset. Fungal sequences were pairwise aligned to generate a distance matrix, which was clustered into OTUs using the average neighbor algorithm (UPGMA) at a 97% similarity threshold. Low abundance fungal OTUs represented by fewer than ten sequences were removed from the dataset. Finally, NCBI’s Basic Local Alignment Search Tool (BLAST) (https://blast.ncbi.nlm.nih.gov/Blast.cgi) was used to identify any OTUs assigned to unclassified fungi. BLAST revealed 335 non-fungal OTU assignments containing 1,111,222 sequence reads, which were subsequently removed from the dataset prior to downstream analyses. These OTUs belonged to the Kingdoms Animalia, Eubacteria, Plantae, and Protista. The majority of the OTUs (41.2%) were assigned as algae followed by plants (20.5%), protozoa (13.4%), insects (13.1%), uncultured eukaryotes and fish (8.95%), and bacteria (2.68%). Fungal OTUs present in the negative PCR or dissection controls (157 OTUs accounting for 349,100 sequences) were also removed from the dataset resulting in a final filtered fungal dataset consisting of 3,415 OTUs and 4,259,127 sequences.

### Amplicon data analysis

To account for unequal sequencing depth while retaining rare taxa, we performed all downstream analyses on a modified filtered fungal dataset (described above) that was not rarefied but excluded all paired mosquito samples for which we obtained <1,500 sequences from either the gut and/or carcass (84). This resulted in elimination of all mosquito gut and carcass samples from site 3, two gut and carcass pairs from site 6, and a single gut and carcass pair from sites 2, 4, and 7 (**Table S5**). Fungal diversity and community composition analyses were conducted using mothur (v.1.38.1) (80). Nonparametric Wilcoxon signed-rank tests were then used to compare alpha diversity (OTU richness, Chao1, and Shannon’s H diversity indices) in the water and mosquito gut and carcass samples.

To compare the mycobiota composition in the same samples, we computed pairwise Bray-Curtis dissimilarity matrix and visualized them by principal coordinates analyses (PCoA) using mothur (v.1.38.1) (80). The compositional differences were inferred via PERMANOVA and MANOVA to determine whether fungal communities clustered based on site or mosquito tissue type (gut vs. carcass). Additional ANOVAs were performed on significant MANOVA factors to determine which axis or axes were responsible for the clustering observed on the PCoA plots. To determine whether the guts or the carcasses were more similar to the breeding water, we performed Wilcoxon matched pairs signed-rank tests comparing the Bray-Curtis distances between the breeding water and larval gut versus those of breeding water and larval carcasses for each larva from a given site. To determine which OTUs may underlie the inferred community differences, we identified indicator OTUs that were disproportionately abundant in either mosquito guts or carcasses using the indicator value (IndVal) method (85). In addition, we determined the degree of preference of OTUs for mosquito guts or carcasses using the point-biserial correlation coefficient (rpb) (86). IndVal and rpb analyses were performed using the Indicspecies package implemented in R (87). For both IndVal and rpb analyses, 9,999 iterations were used to determine whether the OTUs were significantly associated with mosquito guts or carcasses. FUNGuild database (88) was used to determine the ecological role of the fungal OTUs present in the heat map. Fungal OTUs were assigned to six ecological roles: animal pathogenic, ectomycorrhizal, endophytic, epiphytic, fungal parasitic, plant pathogenic, and saprophytic. Although the saprophytic OTUs were divided into dung saprophytic, litter saprophytic, plant saprophytic, soil saprophytic, and wood saprophytic, in our analyses, we assigned all saprophytic OTUs as saprophytes. Wilcoxon signed-rank tests were performed using the GraphPad Prism version 8.4.3 for Windows (GraphPad Software, San Diego, CA, USA), while MANOVAs and ANOVAs were performed using R (http://www.r-project.org/).

## Data availability

Paired sequence data (.fastq files) are available in the National Center for Biotechnology Information (NCBI) Sequence Read Archive under BioProject number PRJNA634912.

## Acknowledgments

We thank Samantha Fox, Kansas State University, for providing us input on sample library preparation. In addition, we thank Jordan Block, Kansas State University, for helping us retrieve literature of the fungal species identified in our study. This study was supported by funding from the National Institutes of Health grant number R01AI140760, the USDA-ARS Specific Cooperative Agreement 58-5430-4-022, and the USDA National Institute of Food and Agriculture, Hatch project 1021223. Support to KLC was provided by USDA NIFA (2018-67012-28009), NSF (IOS-2019368), and the University of Wisconsin-Madison. This is contribution no. 20-310-J from the Kansas Agricultural Experiment Station. Its contents are solely the responsibility of the authors and do not necessarily represent the official views of the funding agencies.

